# Terminus enables the discovery of data-driven, robust transcript groups from RNA-seq data

**DOI:** 10.1101/2020.04.07.029967

**Authors:** Hirak Sarkar, Avi Srivastava, Héctor Corrada Bravo, Michael I. Love, Rob Patro

## Abstract

**Motivation:** Advances in sequencing technology, inference algorithms and differential testing methodology have enabled transcript-level analysis of RNA-seq data. Yet, the inherent inferential uncertainty in transcript-level abundance estimation, even among the most accurate approaches, means that robust transcript-level analysis often remains a challenge. Conversely, gene-level analysis remains a common and robust approach for understanding RNA-seq data, but it coarsens the resulting analysis to the level of genes, even if the data strongly support specific transcript-level effects.

**Results:** We introduce a new data-driven approach for grouping together transcripts in an experiment based on their inferential uncertainty. Transcripts that share large numbers of ambiguously-mapping fragments with other transcripts, in complex patterns, often cannot have their abundances confidently estimated. Yet, the total transcriptional output of that group of transcripts will have greatly-reduced inferential uncertainty, thus allowing more robust and confident downstream analysis. Our approach, implemented in the tool terminus, groups together transcripts in a data-driven manner allowing transcript-level analysis where it can be confidently supported, and deriving transcriptional groups where the inferential uncertainty is too high to support a transcript-level result.

**Availability:** Terminus is implemented in Rust, and is freely-available and open-source. It can be obtained from https://github.com/COMBINE-lab/Terminus.

**Supplementary information:** Supplementary data are available at *Bioinformatics* online.

## 1 Introduction

RNA sequencing has become the de-facto standard for analyzing transcriptomes, and has found myriad applications from differential expression analysis, to the discovery and assembly of rare isoforms. Despite its widespread use, transcriptome analysis via RNA-seq poses a number of computational challenges. For example, reliably mapping or aligning short RNA-seq reads to the reference transcriptome and quantifying the abundance of transcripts is central to most typical RNA-seq analyses. Due to the nature of shared sequences in the reference transcriptome and genome, the simple problem of finding the locus of origin for a particular read sequence can be quite difficult. The complexity results both from alternative splicing, where the isoforms of a single gene can share multiple identical exons, and from very similar sequences arising in families of related genes. These challenges make the read sequence alone insufficient to determine the origin of the sequencing read. In such a scenario, a single read often maps equally well to multiple reference sequences. This ambiguity in determining the exact target sequence propagates directly to the process of quantification, where it becomes hard to determine transcript-level expression when there is insufficient evidence to choose some transcripts over others as the true origin of sequencing reads.

To tackle these challenges, there has been tremendous growth in the space of computational tools that can effectively align (Dobin *et al*., 2013; Langmead and Salzberg, 2012; Kim *et al*., 2015) short RNA-seq reads to transcriptome and tools that can quantify (Li and Dewey, 2011; Turro *et al*., 2011;Glaus *et al*., 2012; Patro *et al*., 2014, 2017; Dao *et al*., 2014) the abundance of transcripts. However, the inherent uncertainty in transcript abundance that results from ambiguous fragment alignment, even after attempting to model this uncertainty in either a maximum likelihood or Bayesian estimation framework, makes it difficult — and in some cases impossible — to provide a single accurate estimate for the number of reads originating from a specific transcript in a given sample. Gibbs sampling is a useful technique for estimating marginal or joint statistics of complex posterior distributions, and has been used by *mmseq* (Turro *et al*., 2011), BitSeq (Glaus *et al*., 2012), RSEM (Li and Dewey, 2011) and *Salmon* (Patro *et al*., 2017) in uncertainty quantification in expression estimation. This provides downstream tools with the ability to analyze the full posterior, instead of just providing a point estimate.

Furthermore, different downstream tools such as mmdiff (Turro *et al*., 2014), IsoDE (Al Seesi *et al*., 2014), sleuth (Pimentel *et al*., 2017) and swish (Zhu *et al*., 2019) make use of these estimates in order to estimate differentially expressed transcripts or genes with higher accuracy and robustness. While taking samples from the posterior probability distribution provides some insight about the validity of the point estimates for transcript abundances, and this uncertainty can be propagated for the purposes of differential testing, it is often possible for a particular transcript to not exhibit expression in the point estimate altogether, in which case it becomes invisible to the analysis tools. Such cases were noted by Turro *et al.* (2014), who further suggested grouping together transcripts whose abundance could not be confidently estimated into transcriptional groups. Specifically, it was demonstrated that there exist numerous cases where the abundance of an individual isoform cannot be reliably estimated, but the abundance of a small group of related isoforms can be determined accurately and robustly.

In a distinct context, but as a result of the same underlying cause of fundamentally multimapping reads, Robert and Watson (2015) note that these difficulties in mapping can lead to errors in quantification that affect genes of relevance to human disease. Crucially, they highlight that this issue occurs even at the level of genes, and is of concern even if one is not performing a transcript-level analysis. In addition to describing this issue, they identify specific groups of disease-related 958 genes that are affected by this problem, and suggest a gene-level analysis approach whereby groups of genes that share multimapping reads are treated jointly for the purposes of expression estimation and differential analysis. While this approach is quite robust, it is also very conservative, since it precludes transcript-level analysis altogether. Furthermore, depending on the set of all fragments sequenced in a sample, it may still be possible to confidently assess the abundance of a gene, or even a single transcript, even if it shares a large number of mulitmapping reads with other sequenced targets. There may then be utility in directly examining the posterior distributions of abundance estimates to determine when multimapping leads to a high degree of uncertainty in estimating expression, and when, despite the presence of multimapping reads, the abundance of a transcriptional target can be confidently assessed.

The mmcollapse tool (Turro *et al*., 2014) exploits the posterior samples generated by mmseq (Turro *et al*., 2011) to identify transcripts with highly anti-correlated posterior distributions. Some of the transcripts in these groups would otherwise not be properly estimated (would have estimated abundances of 0), or would have such variable posterior estimates that they could neither be quantified confidently nor meaningfully tested for differential expression. However, when the transcripts are treated as inferential groups among the experimental samples, they can be robustly quantified and the group can be assessed for differential expression.

One of the major caveats of this approach is the particular choice of summary statistics used in order to identify similar groups. Specifically, *mmcollapse* attempts to group transcripts such that the minimum pairwise correlation is not too low. While this is a useful feature to assess, we find that it does not always accord with intuition about which transcripts should be grouped as it does not specifically account for inferential uncertainty would be reduced by grouping a specific pair of transcripts. Furthermore, the approach taken by *mmcollapse* is both extremely memory intensive (as all posterior samples, for all expressed transcripts, and for all samples in the experiment, must be held in memory simultaneously) and quite time consuming, as the posterior correlations are computed in every iteration of the algorithm, even among completely unrelated transcripts that have little chance of producing promising candidates for grouping. Finally, *mmcollapse* is, arguably, overly-conservative in the constraints is places on transcripts that may be considered as candidates for grouping. For example, only transcripts with no uniquely mapping reads are considered as potential candidates for collapse. However, we observe that even transcripts with a few uniquely mapping reads may exhibit a large degree of uncertainty in their quantification estimates depending on the total number of reads mapping to such transcripts and the complexity of the patterns of multimapping with related transcripts.

Terminus, the tool presented in the current paper, attempts to address these shortcomings. It takes motivation from *mmcollapse* (Turro *et al*., 2014) as well as from the method proposed in “Surface simplification using quadric error metrics” (Garland and Heckbert, 1997), a notable work in the field of computer graphics, in which densely-tessellated shapes are simplified by approximating (coarsening) the mesh that represents the object. In “Surface simplification using quadric error metrics”, Garland and Heckbert (1997) argue that one way of achieving a visually-appealing approximation is to start with the equivalent network of the visual model and repeatedly contract edges of the network in a manner that leads to minimal visual distortion of the overall shape.

In the same spirit, terminus, reformulates the problem of discovering meaningful inferential groups as a graph simplification problem in which the (sparse) graph that defines what transcripts should be considered as candidates for collapsing is constrained by the read multimapping and conditional probability structure conveyed via the range-factorized equivalence classes (Zakeri *et al*., 2017) produced by *Salmon.* This avoids the need to even *consider* the vast majority of possible collapses. Further, terminus uses the reduction in *inferential relative variance* (Zhu *et al*., 2019) — the reduction in inferential uncertainty that would result by grouping together pairs of transcripts — directly as a metric for optimization. We show that this approach is extremely computationally efficient, and that it leads to groups of transcripts that are both biologically and inferentially meaningful. In complex transcriptomes (like human or mouse), our approach reduces the memory requirement by over two orders of magnitude compared to mmcollapse, and is simultaneously two orders of magnitude faster. We validate our results in both simulated and experimental datasets, and present time and memory benchmarks for running these tools.

## 2 Methods

Numerous quantification tools, including *mmseq* and *Salmon*, encode the structure of mapping ambiguity in the form of equivalence classes. Terminus makes use of a collection of *range-factorized* equivalence classes (Zakeri *et al*., 2017) obtained from *Salmon.*

The overview of the mathematical model for *Salmon* is outlined in Supplementary Section 1. Here we would discuss the data structures that are relevant for terminus. Given a set of transcripts 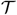 and a set of read sequences 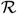, a set of equivalence classes 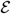, is defined as a function from the domain of set of transcripts 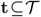 to a natural number denoting the number of reads that are mapped to that group of transcripts, formally 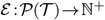, where, 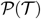 is a power set of all transcripts. In practice the size of the domain of 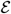 is much smaller than 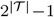. Furthermore *Salmon* extends the notion of naive equivalence classes by adding the measure of mapping quality. Range-factorized equivalence classes can be defined as, 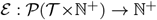. In effect each equivalence class consists of a set of pairs (*t_i_,w_i_*) denoting transcript *t_i_* and a number *w_i_* ∈ (0,1] representing the average conditional probability with which the fragments in this equivalence class arose from transcript *t_i_*. Depending on the granularity with which the range-factorized equivalence classes are defined, the equivalence relation between fragments is determined by the bin, within the range of conditional probabilities, into which they fall with respect to each transcript to which they map. Complete details of how this relation is defined can be found in Zakeri *et al.* (2017). Along with these preliminary structures, *Salmon* also computes Gibbs chains 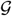, providing samples from the posterior distribution of the model. To be precise, each element 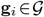 contains the estimated fragment counts for transcript *i*, taken over all (possibly thinned) iterations of Gibbs sampling.

Our goal is to use the range-factorized equivalence classes 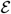 and the posterior samples 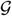 to determine groups of transcripts that exhibit high inferential uncertainty, and then to collect them together into robust transcriptional groups for which the posterior uncertainty is considerably lower. To avoid the exponential space of possible groups, we use the structure over transcripts induced by 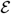 to guide our search, and further restrict each iteration of the algorithm to perform a single pairwise collapse.

First, we collapse transcripts that appear in exactly the same set of equivalence class labels and that have near-identical conditional probability vectors. These are transcripts for which, even without examining the posterior samples, it is clear that no inference algorithm will have sufficient information to tell apart. Specifically, we accomplish this collapse using a partition refinement algorithm (Paige and Tarjan, 1987) where all transcripts start out within a single partition *P*_0_. We then iterate over 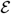 and determine the partitions that should be induced with respect to the current equivalence class 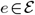. This is simply the subsets of transcripts that have nearly-identical conditional probabilities with respect to *e.* For each such subset *r* in *e*, we refine the current partitioning *P_i_* into *P*_*i*+1_ by replacing each set *S_j_* in *P_i_* that contain elements from *r* with *S_j_* ⋃_*r*_ and *S_j_*\*r.* This process is performed iteratively until we have processed all of the equivalence classes in 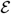.

Next, we define a graph *F* = (*V,E*), constructed over 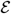 designed to encode the likely candidate transcripts for grouping. We define *V* to be the set of transcripts quantified, where {*υ_i_, υ_j_*} is an edge in *E* if the transcript *υ_i_* and *υ_j_* co-occur in some equivalence class, and either of indication function *h*(*υ_i_*) or *h*(*υ_j_*) is true, and *s*(*υ_i_,υ_j_*) ≤ *τ.* Here *h*(·) is an indicator function that determines if a transcript is, when considered in isolation, a good candidate for possible grouping. We define *h*(·) as,

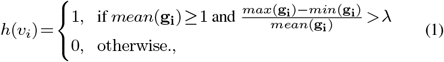

where λ is a user-defined parameter set to 0.1 by default. The score function *s*(*υ_i_,υ_j_*) ≤ *τ* is designed to measure the improvement (decrease) in inferential uncertainty obtained by grouping *υ_i_* and *υ_j_* together. We define

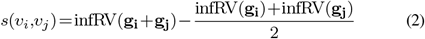

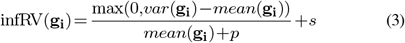

Where *p* is a pseudocount (we use 5 in terminus) and *s* is a small global shift (we use 0.01 in terminus). The infRV(·) function measures the inferential relative variance, and the definition is taken from Zhu *et al.* (2019). The motivation for dividing by the mean is to stabilize the quantification of uncertainty across transcripts with low or high expression. Here, a negative value of *s*(·,·) indicates that, when we sum the posterior samples of the two transcriptional units, the inferential relative variance is less than the average of the inferential relative variance of the individual units. When the value of *s*(·,·) is sufficiently low (see Fig. 1a), then grouping these units together results in a substantial reduction in uncertainty by treating the pair of transcriptional units as a single group. A specific example of such collapse for two transcripts from human is shown in Fig. 2. In practice, we set the threshold for grouping (*τ*) in a data-driven manner by constructing a “background” distribution over the values resulting from evaluating *s*(·,·) on a large number of randomly-selected, expressed transcripts. We note that almost all pairs are not good candidates for collapsing, and so these random draws allow us to approximate sampling from the null distribution of scores for transcripts that should not be grouped. We call this background distribution 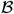.

**Fig. 1:**
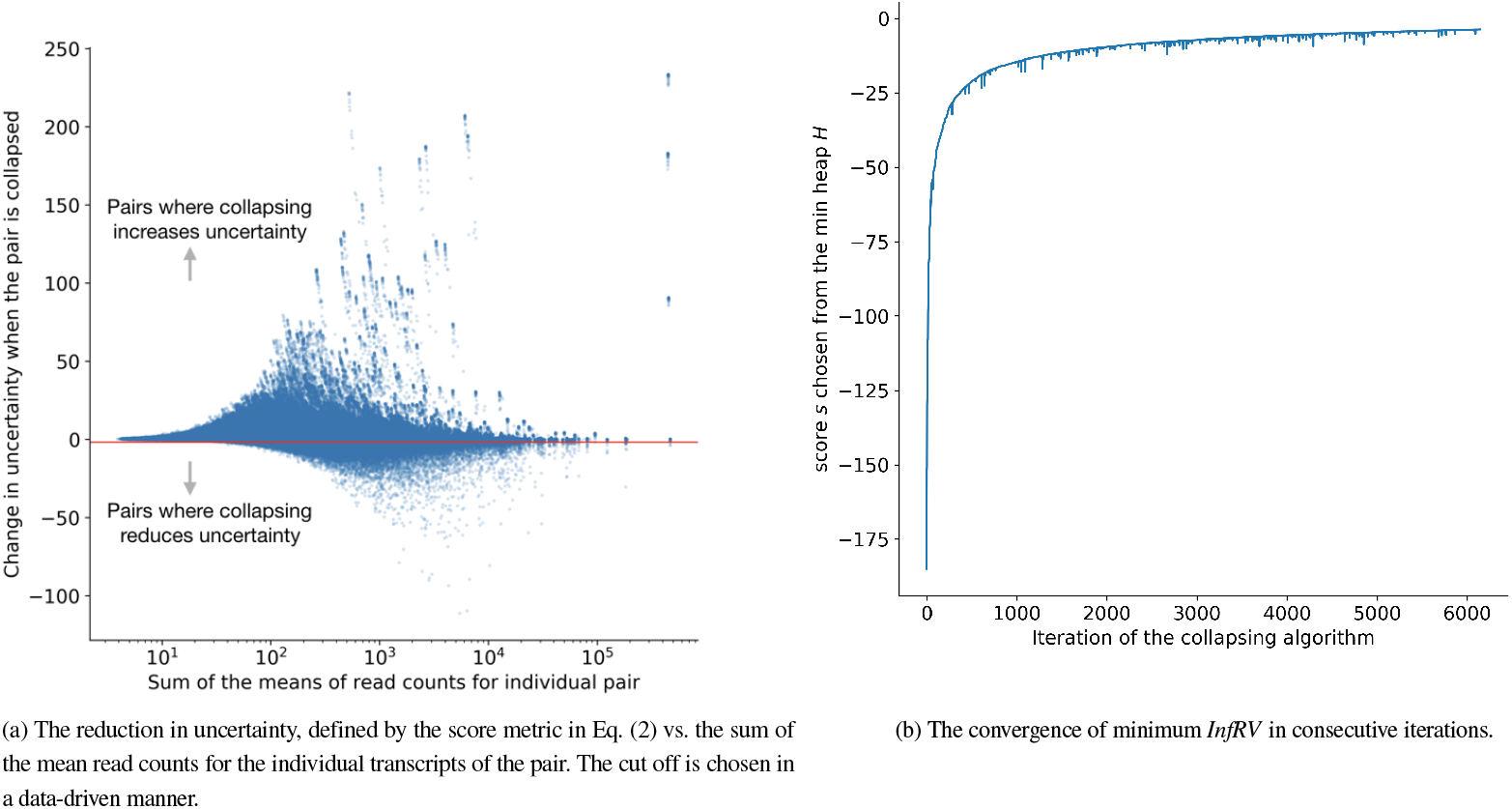
The inferential gain vs the mean of the candidate pairs and the convergence of terminus algorithm.

**Fig. 2:**
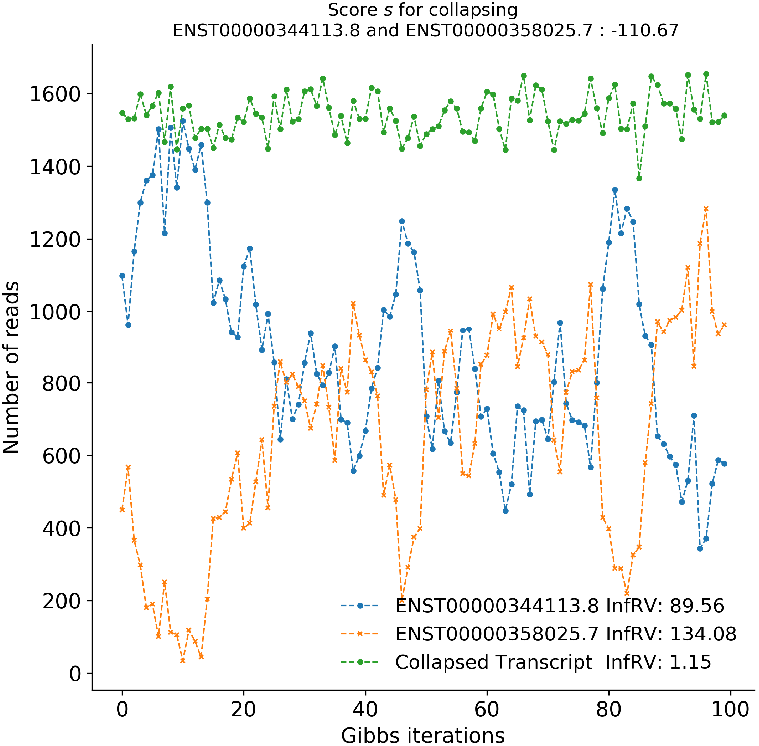
Demonstration of uncertainty reduction by collapsing. Individual transcripts ENST00000310053.9 and ENST00000465259.5 are collapsed, and the corresponding *InfRV* is also reduced.

**Fig. 3:**
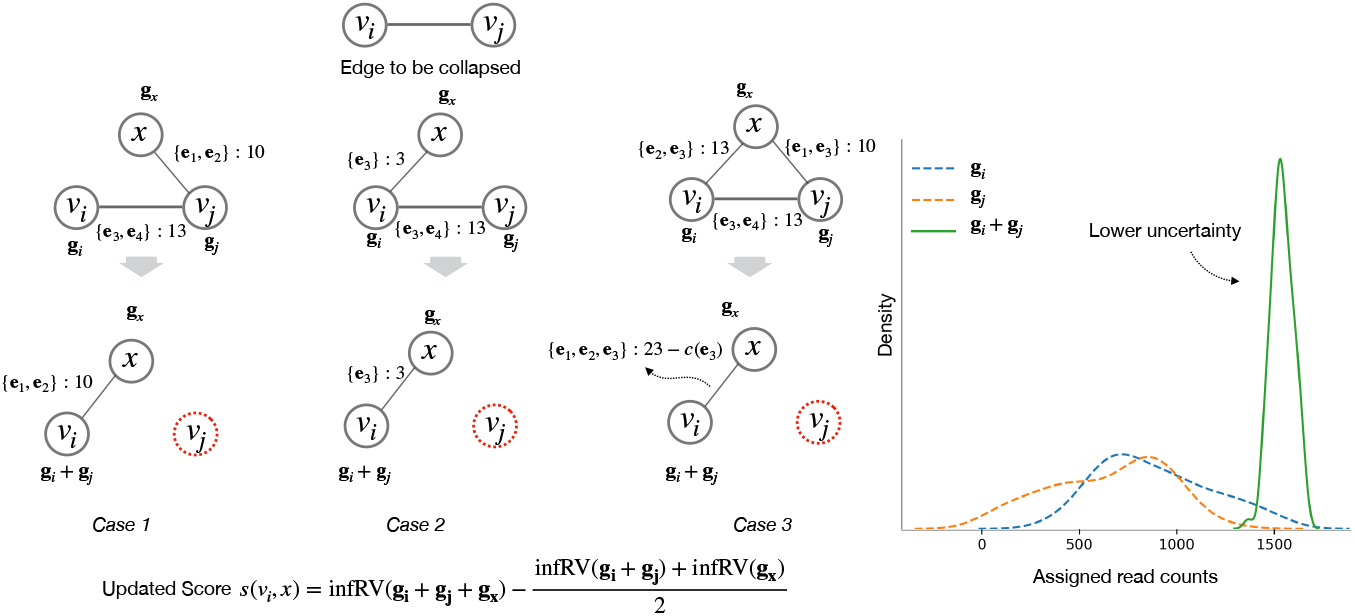
Different possible scenarios that can arise while collapsing a node in Graph F. In case 1 and 2, as the edge property either remains same or transfered to the new edge. In case 3, the construction of edge explained in 2 handles the problem of over counting already counted equivalence class (subtracting count of equivalence class *e*_3_,*c*(*e*_3_)). The right most plot shows the distribution of the individual gibbs samples in dotted line along with the gibbs samples of the collapsed group with much lower uncertainty.

Given a set of transcripts 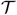, the possible number of transcript pairs to consider for collapse can be 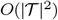. For an organism with a large number of annotated transcripts (such as human or mouse) it is not computationally feasible to enumerate over such a large distribution. Therefore, for choosing a desirable threshold *τ*, we use an iterative sampling approach described in algorithm 2. Specifically, we exponentially increase the number of samples we draw until the resulting threshold *τ* that we would infer changes by less than some small quantity (we use 0.1%). The convergence of algorithm 2 (steps plotted in Supplementary fig. S1) ensures that the final threshold captures a close enough approximation to the true desired value. We set the threshold as a quantile of the background distribution, and choose 2.5% by default. The choice of the percentile is empirical, and chosen so that very few “independent” transcripts might be mistakenly grouped together. This parameter can be used to control sparsity of the graph.

Fig. 1a plots the relation between the scoring function *s*(*i,j*) defined in Eq. (2) and sum of the means of Gibbs samples g_*i*_ and g_*j*_, corresponding to individual transcripts *i* and *j.* The red line shows where the empirical cutoff falls (which changes from one experiment to another). The points below the red line are candidates for grouping.

**Algorithm 1:**
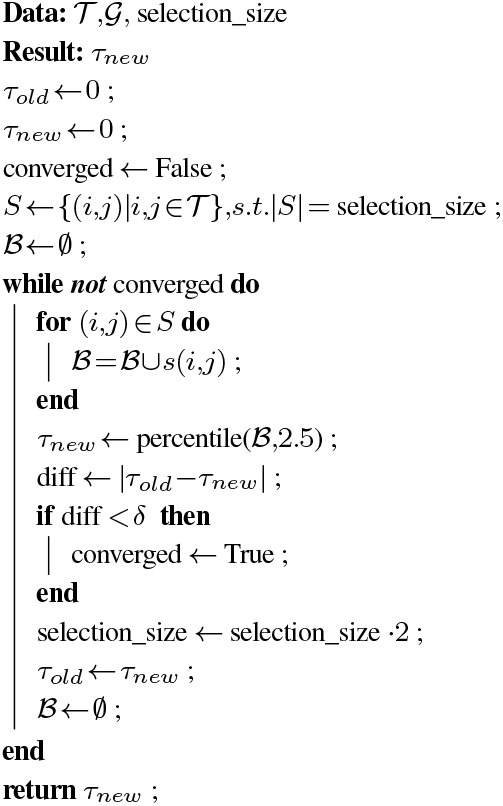
Threshold *τ* selection algorithm.

Given, a graph *F*, a set gibbs samples 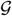, and a threshold *τ*, terminus follows an iterative algorithm for collapsing transcripts. By construction of *F*, an edge {*υ_i_,υ_j_*} ∈ *E* has three different attributes, (i). the score *s*({*υ_i_,υ_j_*}), (ii). a set of equivalence classes *eqlist*({*υ_i_,υ_j_*}) where *υ_i_* and *υ_j_* co-occur and (iii). the total number of reads *c*({*υ_i_,υ_j_*}) that are shared between transcripts corresponding to nodes *υ_i_* and *υ_j_.* Terminus starts off by constructing a min-heap *H* (Cormen *et al*., 2009) over the set of edges *E* where the key for each edge is by the score function evaluated on the vertices sharing this edge. Terminus then iterates over *H* until it becomes empty, at each step collapsing the edge that was popped from the heap.

In each iteration *t*, starting with the current state of graph and the Gibbs samples, 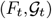, terminus pops an edge {*υ_i_,υ_j_*} from the heap with the minimum score and *collapse* the corresponding end points of the edge and produces a state 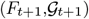. The actual collapse process has a number of cases, but in essence, after collapsing, nodes *υ_i_* and *υ_j_* in the graph *F_t_* becomes a single node in *F*_*t*+1_. Simultaneously, the corresponding vectors g_*i*_ and g_*j*_ from 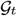 are added to obtain g_*i*_ (to make collapsing more efficient, we associate the collapsed pair with some node *i* from the original endpoints) in 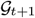.

The collapsing algorithm involves updating the information of the existing edges, and pushing appropriate edges on the heap. At any iteration, given a graph *F_t_* and the edge {*υ_i_,υ_j_*} that has to be collapsed, terminus deletes the edge {*υ_i_,υ_j_*} and includes either of the two nodes, say *υ_i_* (without loss of generality, in practice keeping the smaller numeric index, i.e. *i<j*) in *F*_*t*+1_. Terminus updates 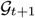 as g_*i*_ ← g_*i*_ +g_*j*_.

The edge set of *F*_*t*+1_ is determined as follows; given the set of adjacent nodes of *υ_i_* and *υ_j_* in *F_t_*, denoted as *adj_t_*(*υ_i_*) and *adjt*(*υ_j_*), any affected node *x* ∈ *adj*(*υ_i_*)⋃*adj*(*υ_j_*) is handled in one of the 3 cases below:

**Case 1:** Given *x* ∈ *adj*(*υ_i_*) and *x* ∈ *adj*(*υ_j_*), an edge {*x,v_i_*} is added to *F*_*t*+1_, terminus recalculates *s*(*x,v_i_*). If *s*(*x,v_i_*) is smaller than *τ*, then the edge with the updated *s*(*x,v_i_*) is pushed into the min heap *H.* The eqlist() and corresponding counts for the pair remain unchanged as this edge already existed in *F_t_.*
**Case 2:** Given *x* ∉ *adj*(*υ_i_*) and *x* ∈ *adj*(*υ_j_*), a new edge {*x,v_i_*} is added to *F_t_*+_1_, terminus calculates *s*(*x,v_i_*). If *s*(*x,v_i_*) is smaller than *τ*, then the edge with the updated *s*(*x,v_i_*) is pushed into the min heap *H.* The eqlist() and corresponding counts for {*x,v_j_*} are copied over to edge (*x,v_i_*), since in the new stated *υ_i_* and *υ_j_* are deemed to be identical.
**Case 3:** Given *x* ∈ *adj*(*υ_i_*) and *x* ∉ *adj*(*υ_j_*), then *x, v_i_* and *υ_j_* forms a triangle in *F_t_.* From all three edges only the edge {*x,v_i_*} is added to *F*_*t*÷1_. Terminus calculates *s*(*x,v_i_*), if *s*(*x,v_i_*) is smaller than *τ*, then the edge with the updated *s*(*x,v_i_*) is pushed into the min heap *H.* The equivalence class list for newly added edge {*x,v_i_*} is recalculated as: *eqlist*({*x,v_i_*}) = eqlist({*x,v_i_*}) ⋃ *eqlist*({*x,v_j_*}). The union of the equivalence class ids saves the edge from over counting those equivalence classes that are shared between all three transcripts *x, v_i_* and *υ_j_.* In the same fashion, the reads that are shared between *x* and *υ_j_* are transferred to the edge {*x,v_i_*}. The shared read counts are calculated as follows

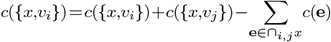 Here ⋂*_i,j_x* signifies a set of shared equivalence classes.

All other nodes and edges remain unchanged in the iteration and are simply “copied over” to *F*_*t*+1_. Similarly, all elements of 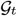 would be copied over to 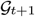 except the changed element g_*i*_ and removal of the element g_*j*_. We observe that the cardinality of *F* and 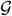 monotonically decreases, ensuring convergence.

Note that throughout the algorithm, the heap maintains the property that any edge currently present in the heap has a score less than empirical threshold *τ.* However, when one endpoint of an edge is modified, the corresponding score of the edge is not directly modified in the heap; such an edge is considered to be “stale.” In order to keep the process of grouping efficient, terminus reuses the same graph in all iterations, keeping state information (i.e. a “last modified” timestamp) encoded as an attribute of the edge. When an edge is updated terminus just updates the corresponding attribute. When an edge is popped from the heap, we first check to ensure that it is not stale before processing the collapse. If the edge is stale, then we recalculate its score and either perform the collapse or re-insert the edge in the heap.

Most typical RNA-seq experiments are comprised of multiple replicates. The process described above, however, is carried out individually per-sample. Terminus takes a two-step approach in order to find coherent groups across the multiple replicates that comprise an experiment. It first groups the individual samples separately, and writes the groups for each sample. This step can be trivially-parallelized. After obtaining individual groups, terminus follows a consensus algorithm in order to find a set of groups to use across all samples.

The consensus procedure starts with individual groups, and constructs a union graph by treating each of the groups as a complete connected undirected graph (clique). For example: given two different groups {*υ_i_,υ_j_,υ_k_*} from one sample and {*υ_i_, v_j_*} from another, terminus first constructs a weighted triangle with end points *υ_i_,υ_j_* and *υ_k_* where each edge has a weight 1. While considering the second group, terminus increases the weight of edge {*υ_i_, v_j_*} by 1. This iterative procedure generates a weighted graph ensuring any edge in the graph represents two transcripts that belong to the same group in at least one sample. Terminus further prunes the union graph and removes the edges that have a weight below a user-defined threshold. The consensus mechanism ensures that a pair of transcripts should at least co-occur in a specified number of samples to qualify for the final grouping. Subsequently the final group that is common for all the samples is extracted by writing the connected components of the pruned graph. Note that these groups are deemed to be universal across samples and replicates. The default consensus threshold is 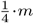 where *m* is the number of samples in the experiment.

### 2.1 Datasets and Evaluation

Different datasets, including both simulated and experimental data and ranging across different organisms were used to demonstrate the benefit of using collapsed groups and the ability of terminus to determine meaningful and robust groups.

#### Simulated dataset on Human

We used polyester Frazee *et al.* (2015)-generated simulated RNA-seq data, curated by Love *et al.* (2018), to assess accurate estimation when the true expression is known. The actual experiment was derived from a joint distribution of mean and dispersion values from GEUVADIS samples (Lappalainen *et al*., 2013). For the current experiment, we have chosen a 4 vs. 4 subset from the original set of 12 vs. 12 samples. The experiments consist of paired end 100 base pair reads (see Love *et al.* (2018) for more detail). We refer to this dataset as *simulated* 4 *vs.* 4 *human* dataset.

#### Simulated dataset with diploid transcriptome from Mouse

The presence of a fully-diploid transcriptome exacerbates the challenges caused by transcript sequence similarity. The mapping uncertainty in such case can greatly impair the quantification estimates. To capture such an extreme scenario, we have produced a diploid transcriptome following the same pipeline described by Raghupathy *et al.* (2018). The diploid transcriptome (named as the N×P transcriptome) combines a cross between NOD/ShiLtK (NOD) and PWK/PhJ (PWK) strains of mice. The final transcriptome is obtained by running `prepare-rsem-reference` on the hybrid `gtf` and reference genome file. The hybrid **gtf** is produced by running `g2gtools` and `emase` (Raghupathy *et al*., 2018). The full script for producing such a transcriptome is provided in the repository. The paired-end read files from N×P transcriptome are generated using polyester with the true counts obtained from running *Salmon* on a real mouse RNA-Seq experiment (accession number `SRR207106`). We refer to this dataset as *simulated allelic mouse* dataset.

#### Experimental RNA-seq samples from Brooks et al. (2011)

The experiment widely known as *pasilla* (Brooks *et al*., 2011) is an ensemble of 6 vs 6 RNA-seq experiment (with NCMI GEO accession numbers GSM461176 to GSM461181) that studies the effect of RNAi knockdown of Pasilla, which is the ortholog in Drosophila melanogaster of NOVA1 and NOVA2 mammallian genes. The same experiment is also used by Turro *et al.* (2014) to demonstrate the effect of grouping.

Given the above mixture of simulated and real experiments, we have run two sets of tools to produce groups, (i) terminus on the output of *Salmon* and (ii) *mmcollapse.* There are different ways to produce input for *mmcollapse.* The *mmcollapse* run is preceded by a run of *mmseq,* which takes BAM files as input. To make the comparison of *mmcollapse* and terminus as consistent as possible, we have used Salmon-produced BAM files for running the *mmcollapse* pipeline. For all experiments, *Salmon* is run with `–numGibbsSamples 100` option in order to generate Gibbs samples. We used the `–hardFilter` parameter for producing the BAM files. In case of hits with different mapping scores, `–hardFilter` keeps only the hits with the best score. This parameter is chosen carefully to follow the equivalent `Bowtie` (Langmead et al., 2009) parameters mentioned in Turro *et al.* (2014) and Turro *et al.* (2011).

Evaluation of the quality of the collapsed groups is inherently a difficult task. Since there can be many possible groupings given a transcriptome dataset, comparing one grouping verses another requires biological validation. For simulated datasets, we validated the results by using the Spearman correlation and mean absolute relative difference (MARD) between the grouped estimates and corresponding true abundances. To be precise a given a grouping *P* is defined as a partition over the set of all transcripts, allowing singleton partitions, denoting the un-grouped transcripts form their own groups. While assessing a particular grouping *P*, we induce the same partitioning over the ground truth abundances. Given a set of true read counts *ρ_true_* = {*ρ*_1_,…,*ρ_N_*} for *N* transcripts and the estimated counts 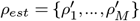 where *M* is the number of groups (including singleton groups), we define a partition *P* induced on *ρ_true_* as,

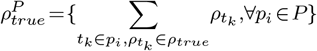

We use 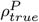 and *ρ_est_* for each of the two tools evaluated in the current manuscript.

## 3 Results

### 3.1 Quantification comparison on simulated data

The two different simulated datasets that are used to demonstrate the utility of the grouping algorithm implemented in terminus pose challenges of distinct natures. While the simulated 4 vs. 4 human dataset is designed to capture aspects of real-world human transcript expression, the simulated allelic dataset from mouse represents the tremendous sequence ambiguity imposed by a diploid transcriptome and its resulting effect on quantification. We demonstrated that, on both the datasets, terminus improves the accuracy of quantification results over *Salmon* and *mmcollapse* under diverse metrics.

The simulated 4 vs. 4 human dataset contains realistic GC bias estimated by alpine (Love *et al*., 2016) from experimental samples from the GEUVADIS study (Lappalainen *et al*., 2013). If these realistic biases are not properly modeled, accurate quantification at the transcript level is impaired. By virtue of the design matrix this experiment simulates differentially expressed transcripts and the nontrivial variability between the samples can also pose challenging problems to the collapsing algorithm. The global Spearman correlation and MARD values presented in Table 1 summarizes those metrics across 8 different samples and takes the average. While *Salmon* itself performs fairly well in this dataset, we observe the groups induced by terminus further improves the accuracy of abundance estimates. The poor performance of *mmcollapse* is due to very noisy low abundance values given to truly unexpressed transcripts. The performance of *mmcollapse* improves substantially when only considering truly expressed transcripts.

**Table 1.** Spearman correlation and MARD (mean absolute relative difference) are calculated with respect to ground truth under for both the simulated 4 vs. 4 human dataset (termed as Human), and the simulated allelic dataset from mouse (termed as Mouse)

| Datasets | Correlation (Spearman) |  |  | MARD |  |  |
| --- | --- | --- | --- | --- | --- | --- |
|  | *Salmon* | *mmcollapse* | terminus | *Salmon* | *mmcollapse* | terminus |
| Human | 0.94 | 0.78 | 0.96 | 0.11 | 0.15 | 0.09 |
| Mouse | 0.91 | 0.81 | 0.95 | 0.12 | 0.12 | 0.07 |

Fig. 4 compares the quantification results at a more granular level under different constraints on the true and estimated counts. Among transcripts that are truly expressed, *mmcollapse* more often mis-assigns reads compared to *Salmon* or terminus. The skewness of histogram from *mmcollapse* suggests that it tends to underestimate the true counts of transcripts. This effect is also visible in the corresponding scatter plot at the top left corner. The spread of the red points signify deviation from the true counts. The scatter plot from terminus *shrinks* these mis-estimates towards the diagonal by putting them into groups improving the overall correlation.

**Fig. 4:**
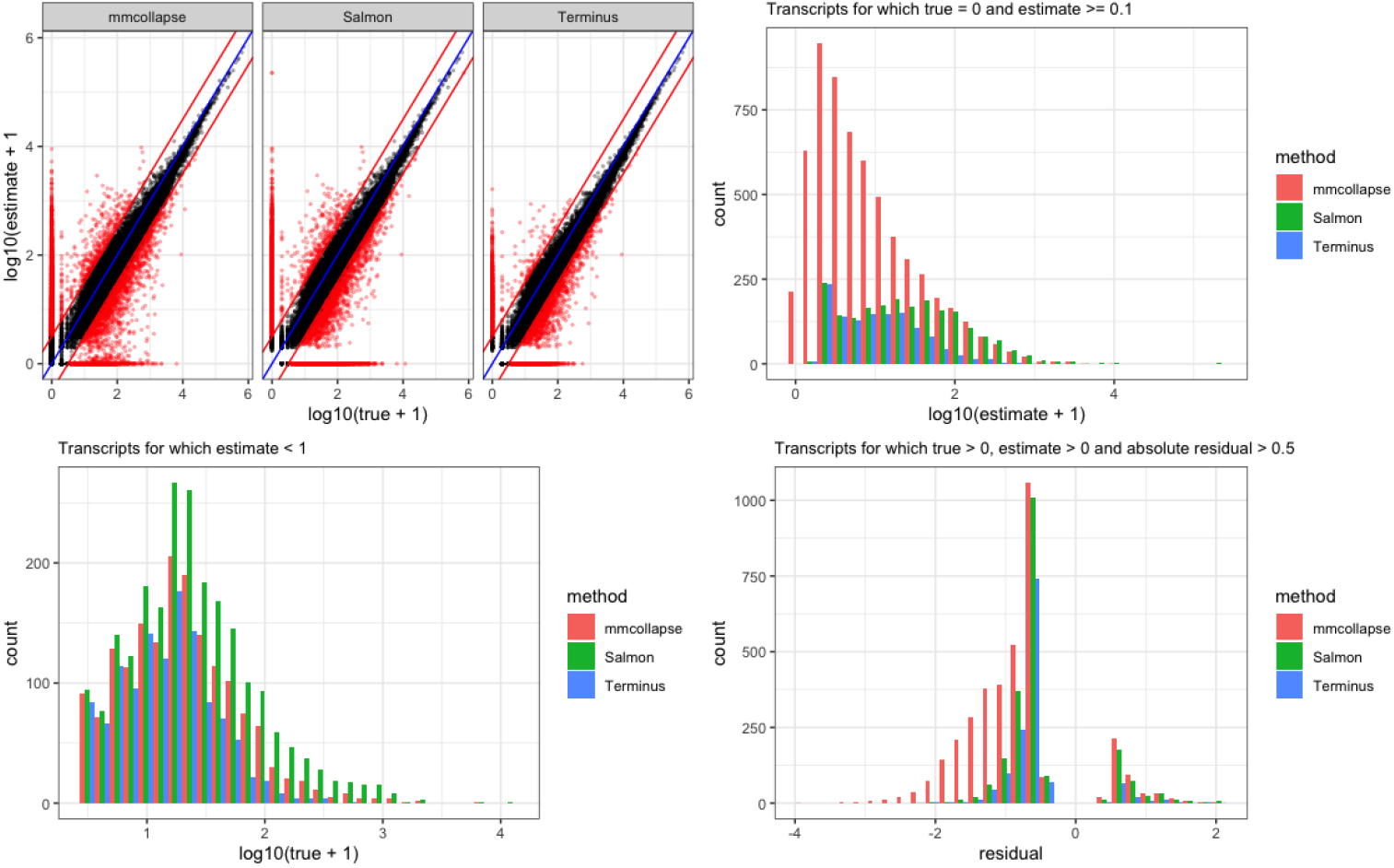
A comparative view of three different tools on the simulated 4 vs. 4 human dataset. The scatter plot on the top right panel asseses the performance of estimates vs truth for three tools *mmcollapse, Salmon* and terminus. The histogram at the top right corner shows the frequency of mis-estimated transcripts which are not expressed in the ground truth. Here only those transcripts for which the tools have at least assigned one read are considered. This helps to get rid of the noise introduced by *mmcollapse* that leads to degenerate values. The histogram at the bottom right shows the frequency of lowly expressed transcripts. The rest of the transcripts (that are expressed) are shown in bottom right plot. Here the x-axis (termed as residual) is the difference between the log transcriformed values of true and estimated expression. Negative residual values signify underestimation while the positive values signify overestimation.

A similar plot for simulated allelic dataset from mouse is shown in Fig. 5. Owing to the diploid transcriptome, we see considerable ambiguity in the underlying sequence. It reduces the overall correlation for all the tools and further affects the convergence of the *mmcollapse* collapsing algorithm. Terminus consistently produces estimated counts closer to truth. The corresponding scatter plot capture the shrinkage of the transcripts groups toward the diagonal which are otherwise mis-estimated by *Salmon.*

**Fig. 5:**
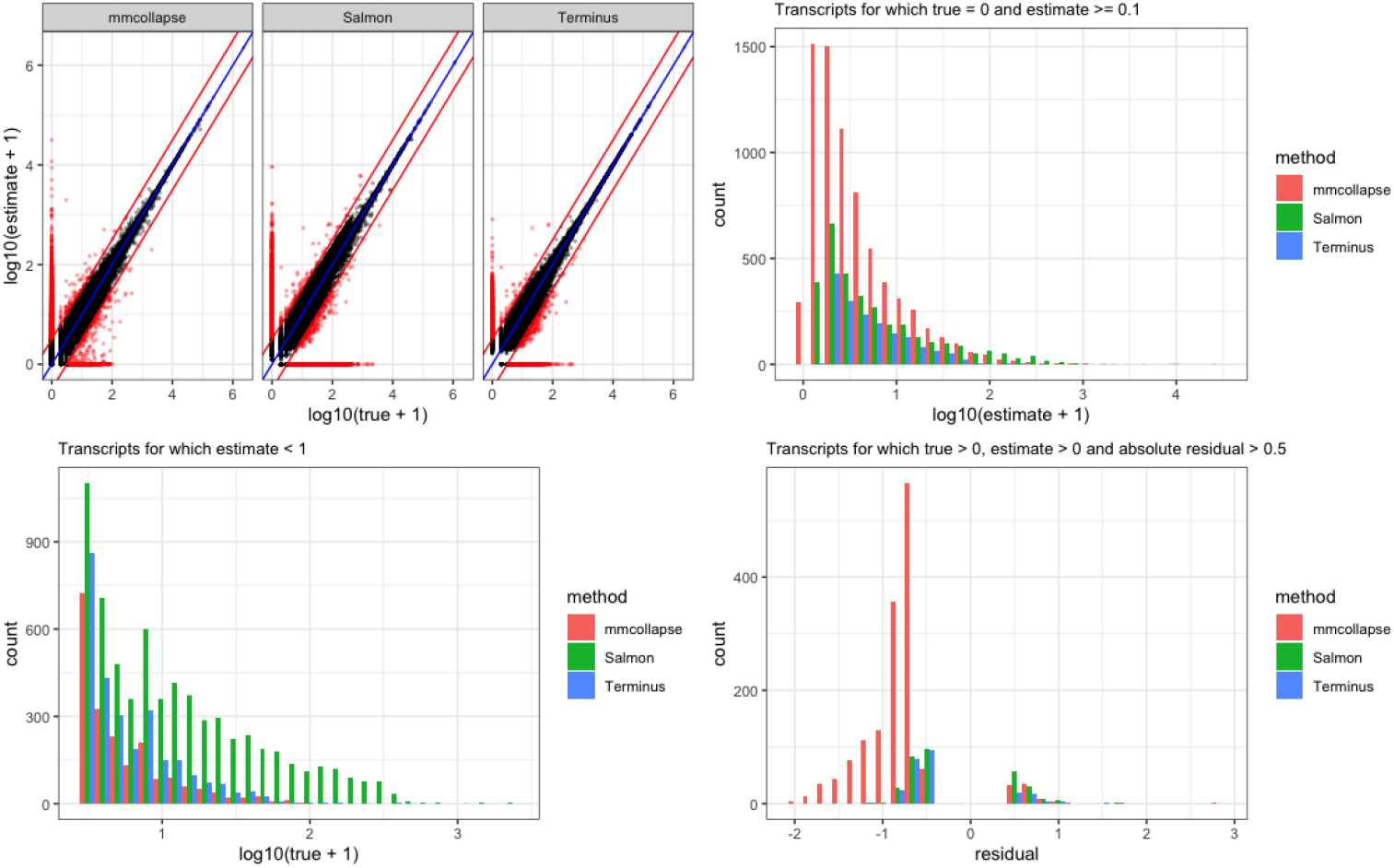
Accuracy of *Salmon* with and without grouping and *mmcollapse* on the simulated allelic dataset is shown in the plot. The metrics are similar to that of Fig. 4

#### Allelic imbalance

Due to presence of highly similar pairs in a diploid transcriptome, it is often a challenging task for a quantification tool to assign reads to the correct allele, especially when both of the alleles are equally likely in terms of the confidence of the alignment. The proportion of two different alleles present in an experiment is often termed as the *allelic imbalance.* When the allelic imbalance is close to 0.5 — when both alleles are expressed equally — accurate estimation becomes particularly difficult. The reason being the fact that there is almost equal prior for assigning a read to either of the candidates. In such cases, the maximum likelihood estimators often prefer one allele over the another, mis-estimating the resulting abundances. We pinpointed such cases by investigating cases where the true allelic imbalance is restricted to an interval of 0.45 to 0.55, focusing on the region where the uncertainly among the alleles is highest.

Supplementary Figure S8 plots the allelic imbalance predicted by *Salmon* (in y-axis ratio of the expression values for individual alleles estimated by *Salmon*) vs. the true allelic imbalance (ratio of true expression of the alleles of a transcript). The color of the point is determined by the fact if the transcript is grouped by terminus (here blue when grouped and orange when not). To get a closer look at the mis-estimation problem in these cases Supplementary Figure S8 zooms the x-axis to the range 0.45 to 0.55. We observe for transcripts with true allelic imbalance 0.5 *Salmon* either over-estimates or under-estimates the true allelic imbalance, resulting in the spread through the the entire y-axis. Meanwhile, the cluster of the points near the very end of *x* = 0.5 vertical line suggests that in those case all the reads are assigned to one of the alleles leading to one allelic imbalance estimates of 0 and 1. As the color (signifying grouped / non-grouped status) suggests, in those cases, terminus correctly identifies the present uncertainty and groups the alleles together.

To quantify the benefit of grouping in the context of allelic imbalance, we have considered the cases of mis-estimated dominant alleles. Specifically, such cases occur when the allelic imbalance ratio is reversed and the allele that truly has higher expression is estimated to have lower expression and vice-versa. We measured the proportion (*r_sal_*) of such cases versus the total number expressed alleles (counting an allele pair only once). We also measured the same metric after grouping. That is, we measured the ratio (*r_term_*) where the numerator is the number of number of mis-estimated dominant alleles that are not grouped, and the denominator is the number of expressed alleles where neither allele from the pairs was grouped. For the simulated allelic dataset we found *r_sal_* to be 0.1 and *r_term_* to be 0.04, signifying the fact that terminus is capable of preferentially grouping together alleles whose allelic ratios are highly uncertain, and where the esimates of *Salmon* are otherwise most-likely to be incorrect.

### 3.2 Quantification on real datasets

The *Pasilla* dataset is an experimental RNA-seq experiment in D. melanogaster. It comprises of both single-end and paired-end RNA-seq reads. Owing to rapid alternative splicing (AS), prevalent in Drosophila, the organism becomes specifically interesting in the context of evaluating uncertainty induced by extensive AS. Almost 20-37% multi-exon genes are alternatively spliced (Gibilisco *et al*., 2016). The isoforms within a gene that share one or multiple long exons are often very hard to distinguish, resulting in low-confidence estimates. As the dataset does not contain the true quantification values, we measured other biological attributes to validate the grouping.

From the 30,597 transcripts, terminus and *mmcollapse* grouped 7904 and 4388 transcripts respectively, distributed in 3025 and 1835 groups. As expected, most of the transcripts in the group originate from the same gene. When groups contain transcripts from multiple genes, these genes belong to either the same gene family or to a closely-related gene family.

Fig. 6 depicts this phenomenon through a histogram plot. In Fig. 6a, the x-axis is the number of genes to which all the transcript within one group can be mapped. For example, for groups that belong to the bar corresponding to *x* = 1, all transcripts come from a single gene; meaning that all the transcripts are isoforms of each other. Similarly, Fig. 6b shows another level of summarization, where the transcripts of a group are mapped back to their gene families. We see in most of the cases that the groups can be mapped back to a small number of gene families. This behavior is expected, as gene families share considerable sequence information and are likely to give rise to related and uncertain expression estimates. Note that *terminus* is agnostic to the annotation of the underlying organism, yet the data driven partitions inherently identify the genes and families annotated in the underlying reference.

**Fig. 6:**
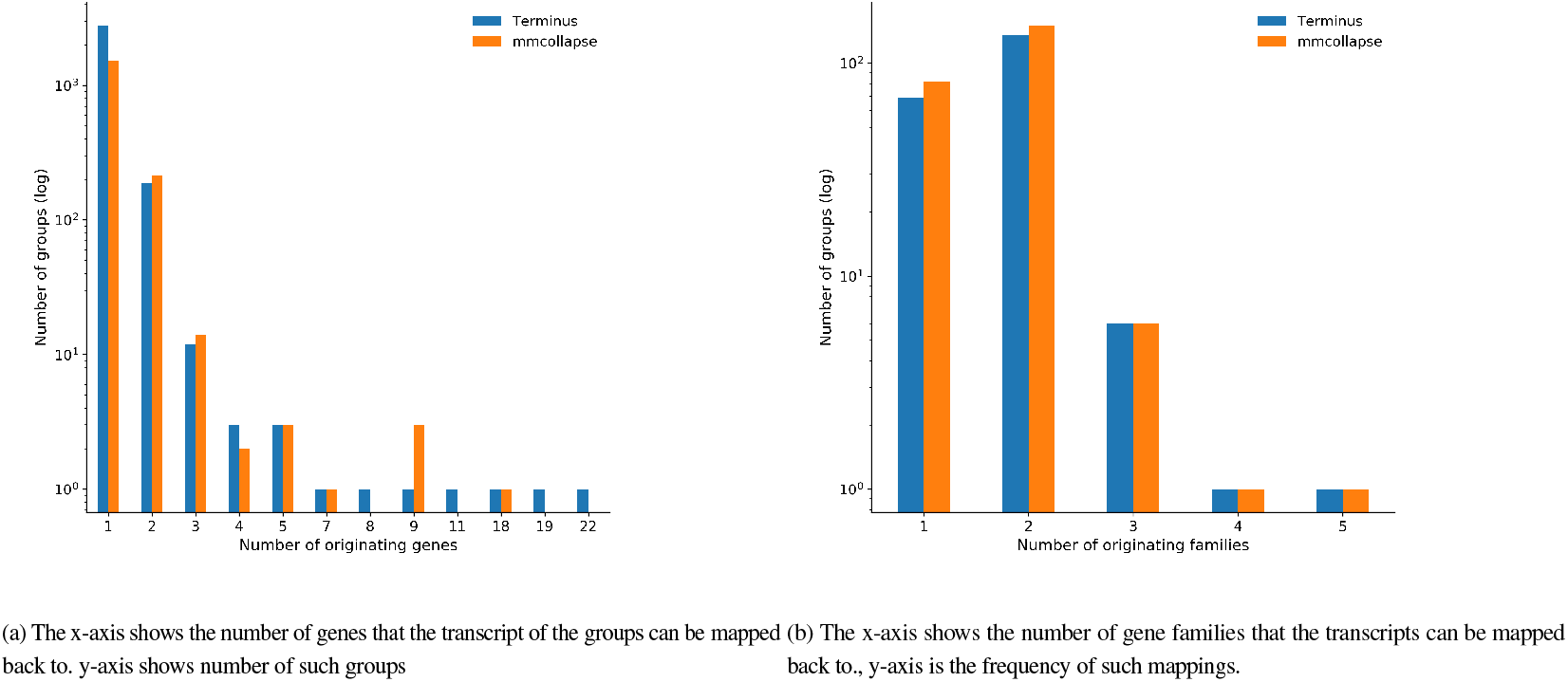
Biological significance of the groups produced by Terminus and *mmcollapse.*

We further observe the group sizes (defined by number of transcripts in a group) generated by terminus (largest one being 54) tend to be larger from that of *mmcollapse* (largest one being 18). Often these large groups also identify gene families that share large numbers of exons. One such example from the groups formed by terminus, consisting 54 transcripts is from the *para* gene and from the same family of parathyroid hormone-related protein. This group of proteins are included in many other groups. Another such group comes from gene *slo* or slowpoke, that regulates the release of a neurotransmitter.

### 3.3 Computational Performance

Table 2 shows the computational performance of terminus vs *mmcollapse.* terminus takes considerably less time to compute the groups and requires much lower memory to run. This enhanced performance derives from two key attributes of terminus. First, terminus does not consider collapsing transcript pairs that do not appear in any equivalence class. This results in the evaluation of many fewer pairs. Second, terminus uses the underlying graph structure induced by the equivalence classes to order and prioritize the collapses, avoiding the need to constantly recompute candidate pairs. We observe that the performance of *mmcollapse,* both in terms of running time and required memory, greatly varies from one sample to another. One possible reason for such behavior could be the variation in the number of non-unique transcripts. While, in the case of the simulated allelic dataset, *mmcollapse* ran for a long time (we terminated the run after 24 hours), for *Pasilla* dataset it finishes relatively quickly. However, we observe that the memory requirement of the tool often made it very difficult to test it with multiple threads. In those cases, the *mmcollapse* run had to be restricted to 1 thread (e.g. on the simulated allelic dataset, running *mmcollapse* with even 1 thread required ~ 214G of RAM). The enormous speed benefits of terminus suggest that it can be easily incorporated as a part of standard lightweight RNA-seq workflow for finding out groups, with very little computational overhead. We note that we have not included the time and memory requirement for *Salmon* and *mmseq* as *mmcollapse* is the tool compared with terminus.

**Table 2.** The table shows construction time and memory requirements for mmcollapse and terminus

| Datasets | Memory(MB) |  | Time (h:m:s) |  |
| --- | --- | --- | --- | --- |
|  | *mmcollapse* | *Terminus* | *mmcollapse* | *Terminus* |
| Human | 370,675 | 2,841 | 3:46:37 | 2:04 |
| Mouse | 225,457 | 485 | 38:55:44 | 0.12 |
| Drosophila | 4,104 | 310 | 8:10 | 0:10 |

## 4 Conclusion and Future Work

The presence of non-unique sequences can greatly affect the accuracy of transcript quantification. Careful analysis of the posterior samples from the underlying probabilistic model not only provides a measure of uncertainty around a point estimate of abundance, but also indicates which groups of transcripts may have abundances that are particularly difficult to distinguish individually but which have estimable abundance as a group. Terminus demonstrates that Gibbs samples can be used to identify groups of transcripts that exhibit high inferential uncertainty on their own, but which exhibit much lower uncertainty as a group. We show how terminus uses the information encoded in range-factorized equivalence classes, readily available after quantification with *Salmon*, to tremendously accelerate the grouping process. Terminus writes the new expression estimates and posterior samples with the group information in the same exact format as that of *Salmon,* enabling any downstream pipeline that accepts a similar format (Zhu *et al*., 2019) to directly run on terminus output.

The groups computed by terminus represent abundance estimates reported at the resolution that is actually supported by the underlying experimental data. In a typical experiment, this is neither at the gene level nor the transcript level. Some transcripts, even from complex, multi-isoform genes, can have their abundances accurately estimated and with low uncertainty, while other transcripts cannot. Rather than pre-defining the resolution at which the analysis will be performed, and subjecting the results to either overwhelming uncertainty or to insufficient biological resolution, terminus allows the determination of transcriptional groups whose abundance can be confidently estimated in a given dataset, and represents, in this sense, a data-driven approach to transcriptome analysis.

Further, we demonstrate that terminus creates biologically relevant groups that reflect the underlying hierarchy of genes and gene families. This shows the potential of terminus to be utilized for applications of data-driven clustering of biological sequences, such as clustering de-novo contigs (where the annotation is not known) or for clustering related strains in metagenomic samples.

From a conceptual perspective, terminus provides a novel approach for grouping complex interactions between biological sequence without having the prior information about the annotation itself. It first prunes the possible space of pairwise collapses by examining the structure induced by the range-factorized equivalence classes, and later, uses a iterative greedy technique to collapse transcripts that locally maximize the objective being optimized (i.e. the reduction in inferential relative variance). It is possible that this generic framework can be extended to other aspects of grouping and clustering such as taxonomic classification.

## References

Al Seesi, S. et al. (2014). Bootstrap-based differential gene expression analysis for RNA-Seq data with and without replicates. In BMC genomics, volume 15, page S2. BioMed Central.

Brooks, A. N. et al. (2011). Conservation of an RNA regulatory map between drosophila and mammals. Genome Research, 21(2), 193–202.

Cormen, T. H. et al. (2009). Introduction to algorithms. MIT press.

Dao, P. et al. (2014). ORMAN: optimal resolution of ambiguous RNA-Seq multimappings in the presence of novel isoforms. Bioinformatics, 30(5), 644–651.

Dobin, A. et al. (2013). STAR: ultrafast universal RNA-seq aligner. Bioinformatics, 29(1), 15–21.

Frazee, A. C. et al. (2015). Polyester: simulating RNA-seq datasets with differential transcript expression. Bioinformatics, 31(17), 2778–2784.

Garland, M. and Heckbert, P. S. (1997). Surface simplification using quadric error metrics. Proceedings of the 24th Annual Conference on Computer Graphics and Interactive Techniques, pages 209–216.

Gibilisco, L. et al. (2016). Alternative splicing within and between drosophila species, sexes, tissues, and developmental stages. PLoS Genetics, 12(12).

Glaus, P. et al. (2012). Identifying differentially expressed transcripts from RNA-seq data with biological variation. Bioinformatics, 28(13), 1721–1728.

Kim, D. et al. (2015). HISAT: a fast spliced aligner with low memory requirements. Nature Methods, 12(4), 357–360.

Langmead, B. and Salzberg, S. L. (2012). Fast gapped-read alignment with Bowtie 2. Nature Methods, 9(4), 357.

Langmead, B. et al. (2009). Ultrafast and memory-efficient alignment of short DNA sequences to the human genome. Genome biology, 10(3), R25.

Lappalainen, T. et al. (2013). Transcriptome and genome sequencing uncovers functional variation in humans. Nature, 501(7468), 506–511.

Li, B. and Dewey, C. N. (2011). RSEM: accurate transcript quantification from RNA-Seq data with or without a reference genome. BMC Bioinformatics, 12(1), 323.

Love, M. I. et al. (2016). Modeling of RNA-seq fragment sequence bias reduces systematic errors in transcript abundance estimation. Nature Biotechnology, 34(12), 1287.

Love, M. I. et al. (2018). Swimming downstream: statistical analysis of differential transcript usage following salmon quantification. F1000Research, 7.

Paige, R. and Tarjan, R. E. (1987). Three partition refinement algorithms. SIAM Journal on Computing, 16(6), 973–989.

Patro, R. et al. (2014). Sailfish enables alignment-free isoform quantification from RNA-seq reads using lightweight algorithms. Nature Biotechnology, 32(5), 462.

Patro, R. et al. (2017). Salmon provides fast and bias-aware quantification of transcript expression. Nature Methods, 14(4), 417.

Pimentel, H. et al. (2017). Differential analysis of RNA-seq incorporating quantification uncertainty. Nature Methods, 14(7), 687.

Raghupathy, N. et al. (2018). Hierarchical analysis of RNA-seq reads improves the accuracy of allele-specific expression. Bioinformatics, 34(13), 2177–2184.

Robert, C. and Watson, M. (2015). Errors in RNA-Seq quantification affect genes of relevance to human disease. Genome Biology, 16(1), 177.

Turro, E. et al. (2011). Haplotype and isoform specific expression estimation using multi-mapping RNA-seq reads. Genome Biology, 12(2), R13.

Turro, E. et al. (2014). Flexible analysis of RNA-seq data using mixed effects models. Bioinformatics, 30(2), 180–188.

Zakeri, M. et al. (2017). Improved data-driven likelihood factorizations for transcript abundance estimation. Bioinformatics, 33(14), i142–i151.

Zhu, A. et al. (2019). Nonparametric expression analysis using inferential replicate counts. Nucleic Acids Research, 47(18), e105–e105.

