## Supplementary Document for "Terminus enables the discovery of data-driven, robust transcript groups from RNA-seq data"

### 1 Model used in *Salmon*

The basic probabilistic model of *Salmon* (Patro *et al.*, 2017) is adapted from the basic generative model proposed by Li *et al.* (2010). Defining the known nucleotide fraction as $\eta$, set of (read) fragments as $\mathcal{F}$, set of transcripts $\mathcal{T}$ and the transcript to fragment assignment binary matrix as $\mathcal{Z}$, then the transcript level quantification aims to solve,

$$\text{argmax}_{\eta}\mathcal{L}(\eta|\mathcal{F},\mathcal{T})=\prod_{j=1}^{N}\sum_{i=1}^{M}Pr\{t_i|\eta\}Pr\{f_j|t_i,z_{ij}=1\}. \tag{1}$$

*Salmon* performs a variational Bayesian optimization (using a combination of a stochastic collapsed variational Bayesian optimization algorithm (Foulds *et al.*, 2013) in an online inference phase and a variational Bayesian expectation maximization algorithm (Nariai *et al.*, 2013) in the offline phase, with the latter phase being performed over range-factorized equivalence classes (Zakeri *et al.*, 2017)). In addition to the point estimates provided derived above, *Salmon* provides the option to draw samples from the posterior distribution of the parameters using a Gibbs sampling procedure. This sampling procedure starts at the optimized parameters, and performs Gibbs sampling according to the equations introduced in Turro *et al.* (2011). This provides, in addition to the point estimates, samples from the posterior distribution that can be used to assess the confidence in each transcript's abundance. Finally, *Salmon* can also output the range-factorized equivalence classes that it uses internally for inference. These act as a reduced representation of the experiment, which are used downstream in terminus. Further details about the operation and features of *Salmon* are explained in Patro *et al.* (2017).

### 2 The computational pipeline for running *mmcollapse* and terminus

The *mmcollapse* tool is specifically tied to *mmseq* as the upstream inference algorithm. It relies on specific estimates computed by *mmseq*. Likewise, terminus runs downstream of *Salmon* and relies on output generated by *Salmon* (e.g. range-factorized equivalence classes) that is not produced by *mmseq*. Thus, there are certain, unavoidable, differences to the inputs of these tools that cannot be avoided. However, to carry out as fair a comparison as possible, we have attempted to minimize controllable differences between the *mmseq* → *mmcollapse* and *Salmon* → terminus pipelines by using *Salmon* consistently for producing the mapping/alignment results that are provided to *mmseq*. *Salmon* is capable of producing both abundance estimates as well as SAM files containing the alignments that it has computed internally. The default *mmcollapse* pipeline specifies the use of `Bowtie` (Langmead *et al.*, 2009). In order to best match the expectations of *mmseq*, we have used `--hardFilter` option to make *Salmon* produced alignments similar to those of `Bowtie`. The same reference index used for producing these alignments is used by *Salmon* to produce the quantification files and Gibbs samples later used by terminus. *mmcollapse* requires the output file from *mmseq*, therefore *Salmon*-produced quantification results could not be used directly with *mmcollapse*. At the end of the pipeline, both *mmcollapse* and terminus produced grouping files that are used for comparing results.

### 3 Terminus-produced groups and the effect on posterior variance

Supplementary Table 1 shows the exact numbers of transcripts that are collapsed and the number of groups that are formed during the collapsing procedure. We observe that for the simulated 4 vs. 4 human dataset dataset, the number of transcripts that are collapsed is considerably higher in terminus compared to *mmcollapse*. The reason for such difference in the number of groups can be caused by the specific grouping

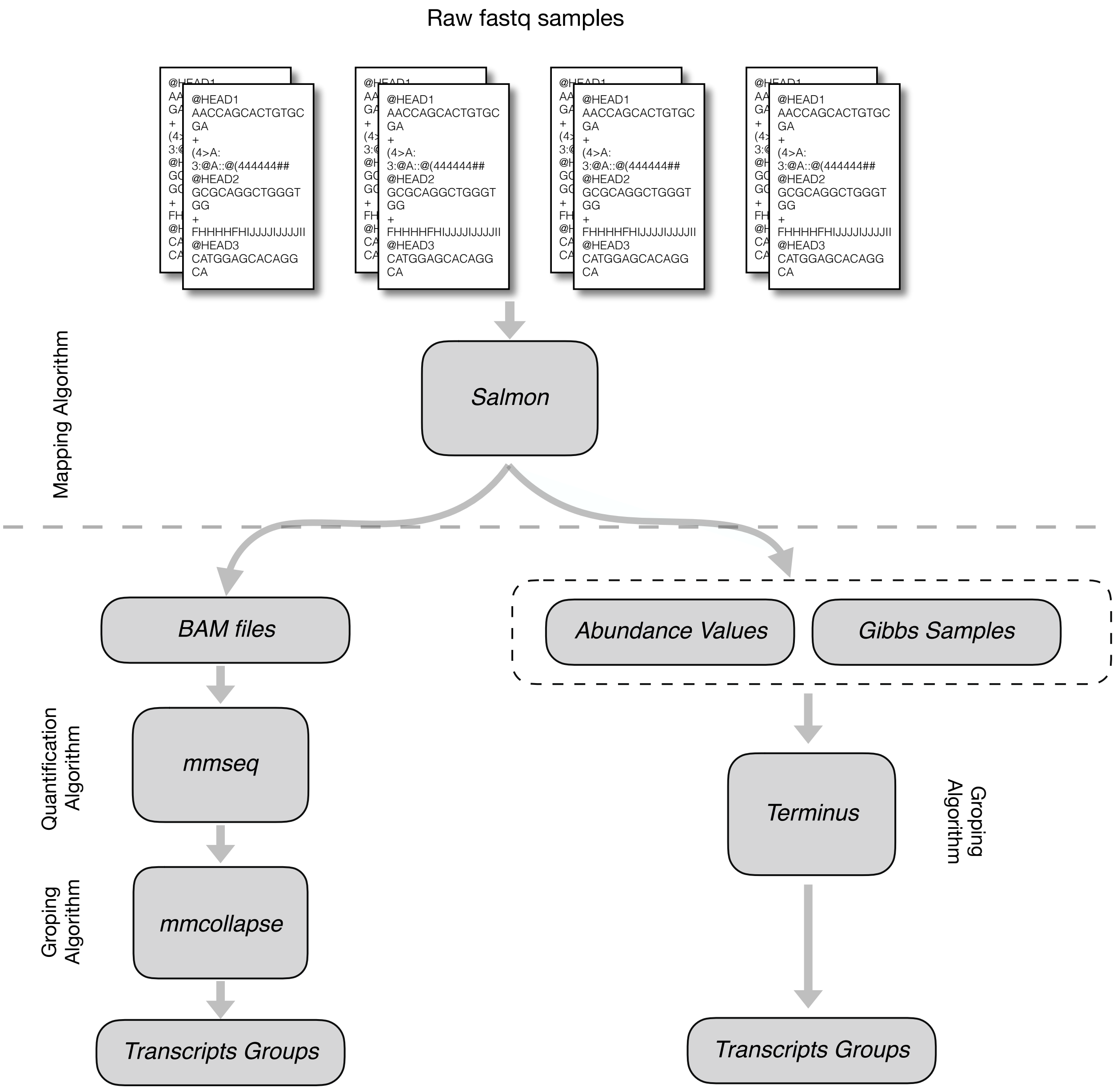


Supplementary Figure S1: The detailed pipeline for running *Salmon* and terminus, while sharing the produced alignments with *mmseq* and *mmcollapse*.

algorithm that *mmcollapse* follows, and the fact that terminus does not *a priori* exclude transcripts from grouping simply because they have some uniquely-mapping fragments. On the simulated allelic dataset dataset, however, the number of groups produced by *mmcollapse* is much higher than that of terminus. On the *Pasilla* dataset dataset, we observe a comparable number of groups formed by both methods.

To further investigate the effect of grouping in one of the samples from simulated 4 vs. 4 human dataset. We have considered all groups with cardinality 2, and measured the change in variance after merging the Gibbs samples from individual candidates. In Supplementary Figure S2 the variance for the merged groups are plotted with respect to the mean of the variances from the Gibbs samples of individual transcripts. We observe that out of 3553 two member groups, 2094 pairs the merged variance is decreased and 1444 cases there is an increase. We further observed while the increase in variance never crosses the difference of 1,

Supplementary Table 1: Group statistics for simulated 4 vs. 4 human dataset (termed as Human), and the simulated allelic dataset from mouse (termed as Mouse) and *Pasilla* dataset (termed as D.Mel)

| Datasets | Number of transcripts grouped | | Number of groups | |
|---|---|---|---|---|
| | terminus | *mmcollapse* | terminus | *mmcollapse* |
| Human | 12623 | 1454 | 4972 | 640 |
| Mouse | 37241 | 53325 | 17554 | 24831 |
| D.Mel | 4863 | 4388 | 2040 | 1835 |

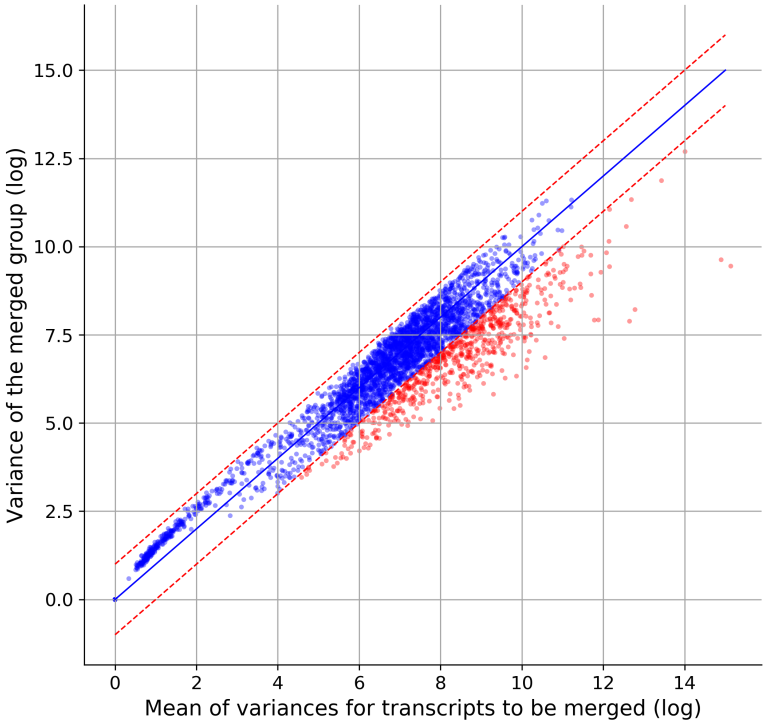


Supplementary Figure S2: The change in posterior variance after merging transcripts compared to the mean variance of the individual transcripts.

the decrease for highly variant values are well beyond that (marked with red color). Noting the *log* scale this positively shows that the terminus produced groups bound the change in variance after the collapse.

#### 4 Detailed comparison between *Salmon* and terminus

In order to highlight the improvement obtained by terminus, we plotted the comparative metrics of *Salmon* and terminus in Supplementary Figure S3 and Supplementary Figure S4 for simulated simulated 4 vs. 4 human dataset and simulated allelic dataset datasets respectively.

Both the datasets categorically show different cases conditioned on the true expression of the tran-

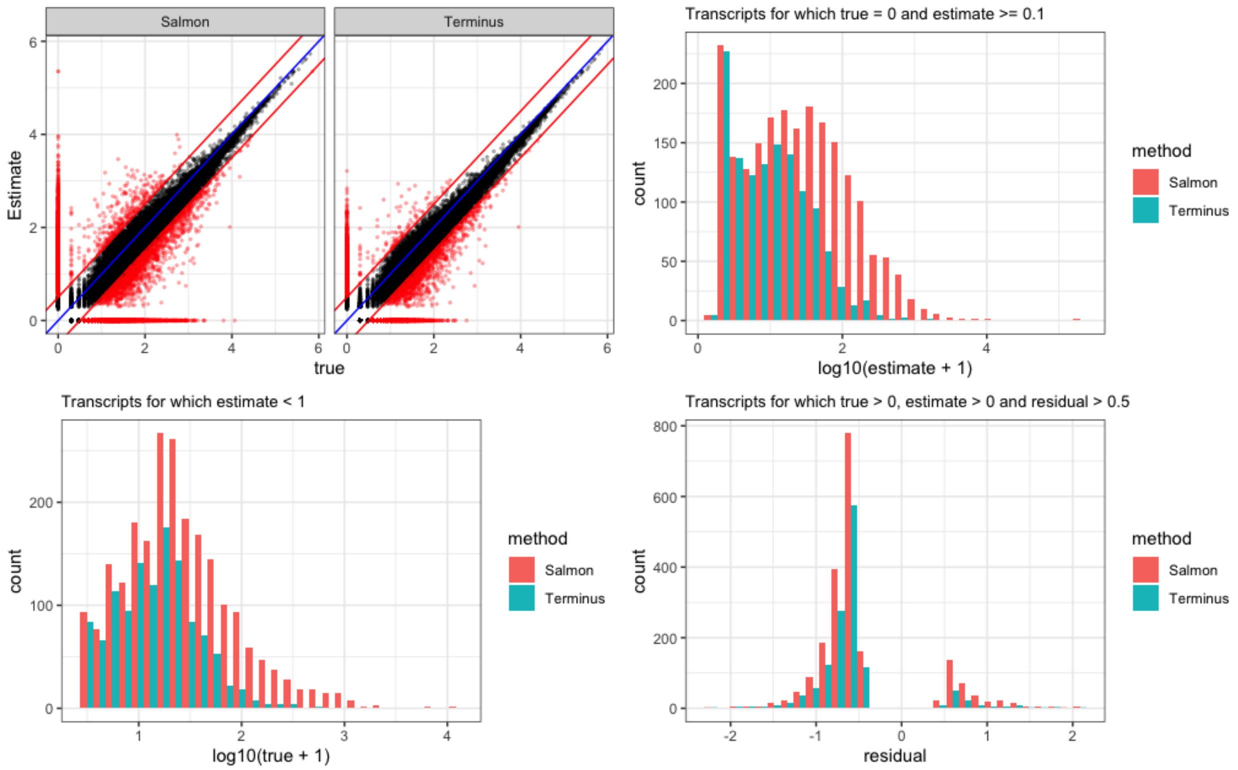


Supplementary Figure S3: A comparative view of *Salmon* and terminus on the simulated 4 vs. 4 human dataset. Among transcripts that are truly expressed. Following the same interpretation as described in Fig. 4, terminus grouping reduces the number of mis-estimated transcripts.

scripts and the corresponding estimates from the respective tools. Starting from the scatter plot at the right top corner, we observe the spread of expressed transcripts that are estimated to be unexpressed by the two tools (the horizontal spread at $y = 0$ marked in red) and the truly unexpressed transcripts that are expressed by the respective tools (the vertical spread at $x = 0$ marked in red). The number of such mis-estimated points are significantly decreased in terminus compared to *Salmon*. Additionally, we observe a shrinkage in the mis-estimated points (points away from $x = y$ line) in the scatter plot for terminus, signifying that the tool has reduced the number of mis-estimated abundances. To capture the magnitude of such mis-estimations in both *Salmon* and terminus, we observe the histograms of the abundances conditioned on true expression values. The first histogram (top-right corner of Supplementary Figure S3 and Supplementary Figure S4) considers the transcripts for which the true expression is zero. The shift of the transcript count distribution for different levels of mis-estimated abundances demonstrates that the magnitude of mis-estimation is more severe in *Salmon*, at the transcript level, compared to terminus, at the group level. The same trend is to be seen for lowly-abundant transcripts (abundance values less than 1 in the bottom left corner). The last plot shows the histogram for transcripts for which $|\log_{10}(y+1)$ - $\log_{10}(x+1)| \geq 0.5$, where $y$ is the estimated abundance and $x$ is the true abundance. In this case also we see that *Salmon* has more mis-estimated abundances than terminus.

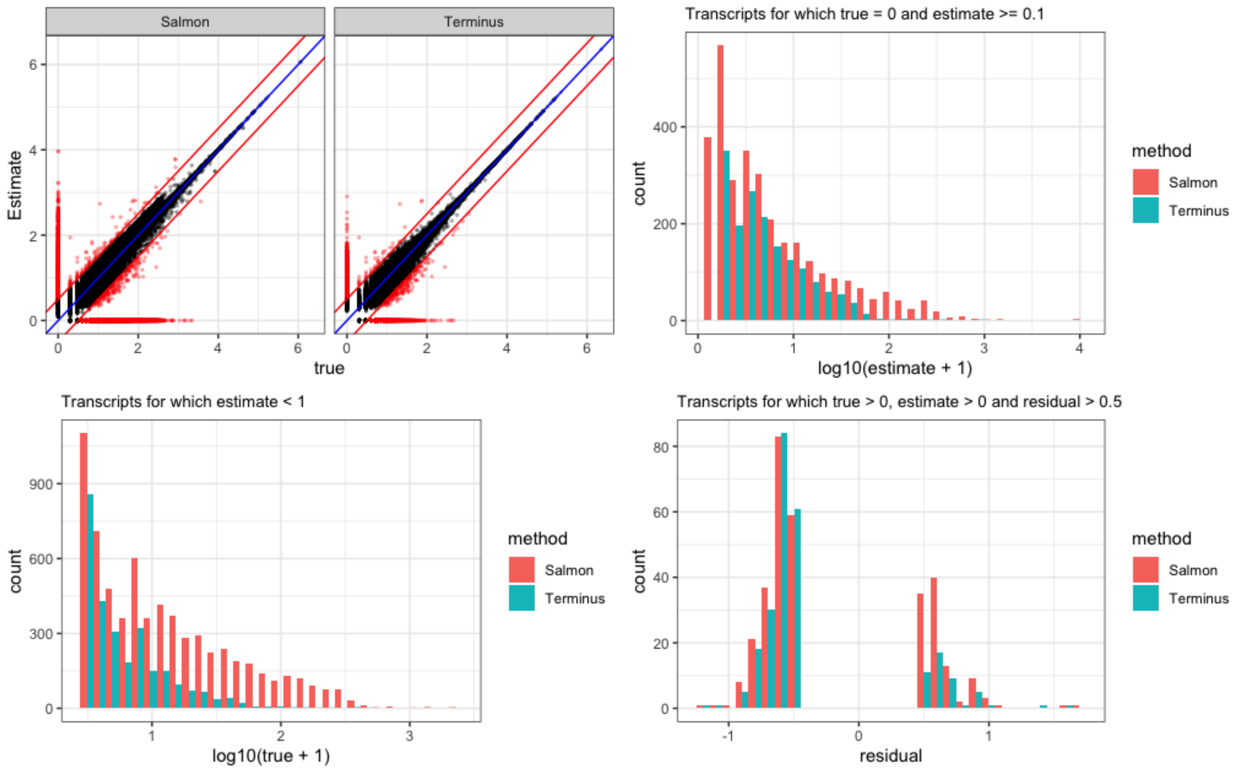


Supplementary Figure S4: A comparative view of *Salmon* and terminus on the simulated allelic dataset. The metrics are similar to that of Fig. 4

#### 4.1 Exploratory analysis for mis-estimated abundances in simulated 4 vs. 4 human dataset

For the simulated 4 vs. 4 human dataset we have selected a few transcripts from the human transcriptome where the transcript abundance estimation from *Salmon* deviates from the simulated counts by a substantial margin. To emphasize the effect of such mis-estimation in the downstream pipeline, we have chosen one of the replicates (among 4) where both the control and treatment samples are taken into account. We compared the log fold change (termed as LFC) of the transcript-level fragment counts simulated by polyester (Frazee *et al.*, 2015) and the counts estimated by *Salmon* between the two samples. Further, we identified only the transcripts for which, i. the LFCs are reversed (i.e. while the true count based log fold change is positive the LFC from *Salmon* counts are negative or vice versa) and ii. the absolute difference of the LFCs are more that 0.5. The goal of such a filter is to consider the transcripts which are estimated to be up-regulated while they are, in reality, down-regulated and vice-versa.

The distribution for the log fold change *for these transcripts with mis-estimated fold changes* is shown in Supplementary Figure S5. We observe that, as expected, the estimated log fold change distribution is different from the true distribution. For this particular experiment, there are 2194 such transcripts. It spans through 232 different gene families. Terminus groups 669 transcripts out of these 2195 into different groups (note that the groups may contain transcripts outside this set).

Supplementary Figure S6 captures this phenomenon of groping graphically. The abundance estimates by *Salmon* are plotted in blue, while group-level abundance estimates (groups of which these transcripts are members) are plotted in red. An arrow originates at a blue point that is to be grouped by terminus, and points to a red point that is the group-level estimate for the group containing this transcript. The

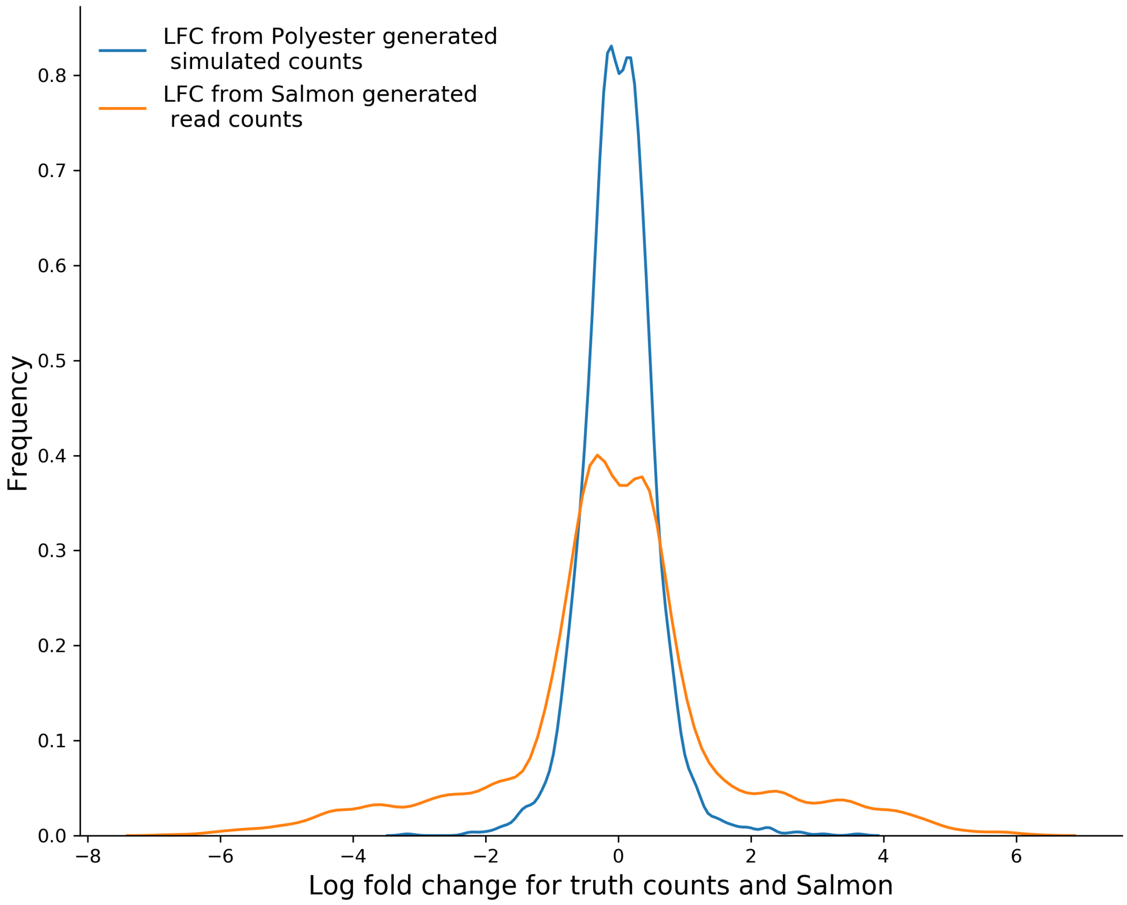


Supplementary Figure S5: Distributions of log fold changes between control and treatment samples from one of the replicates of simulated 4 vs. 4 human dataset. The distribution is on a subset of transcripts as defined in Supplementary Sect. 4.1

pattern of arrows show that the mis-estimated transcripts are away from the $x = y$ line and, when grouped by terminus, the *grouped estimates* are much closer to the $x = y$ line (i.e. the grouped abundances are much closer to the corresponding grouped true counts).

##### 4.2 Exploratory analysis for mis-estimated transcripts in GEUVADIS sample $ERR188204$

We further experimented with a samples from GEUVADIS (Lappalainen *et al.*, 2013), $ERR188204$. Due to the absence of ground-truth, to asses the performance of *Salmon* and terminus, we have created a dataset derived from $ERR188204$ by artificially shortening the reads from the `FASTQ` files. To be specific, 26 nucleotides are trimmed from the 76 nucleotide reads in the original `FASTQ` file. The unaltered dataset is used as the ground truth, while the quantification results on the trimmed dataset are assessed. To further increase the resolution of the performance comparison, we considered transcripts that originates from gene families where there are a numerous transcript isoforms present (the specific selection procedure

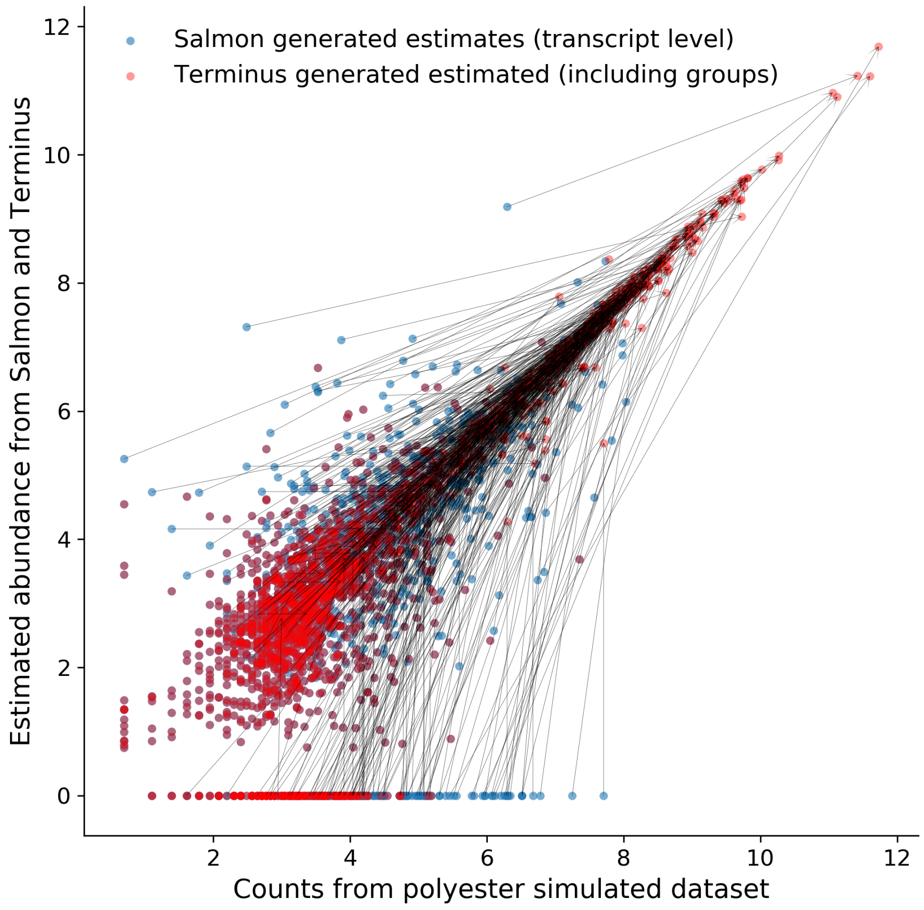


Supplementary Figure S6: The transcripts which are mis-estimated by *Salmon* are often grouped by terminus. The arrows originate from an abundance values estimated by *Salmon* (marked in blue) for a transcript and points to a group (marked in red) that is formed by terminus.

is defined below).

We identify such gene families of interest by calculating the ratio of total number of transcripts versus the total number of genes within that family. The ratio is termed as splicing repertoire (SR). As an illustrative example, the `N-myc downregulated gene family` (NDRG) has 4 genes and 167 transcripts, making it one of the highest SR-scored gene families (with score 167/4 = 41.75). For this experiment, we consider transcripts that satisfies two conditions, namely, i. it belongs to a gene family with SR score more than 10 and ii. it is expressed with a coverage lower than 100 reads. Supplementary Figure S7 (following similar convention as of Supplementary Figure S6) shows the scatter plot for such transcripts from *Salmon* (labeled with blue) and the effect of terminus (red) grouping leading to a shift towards the $x = y$ line, improving the overall correlation considerably.

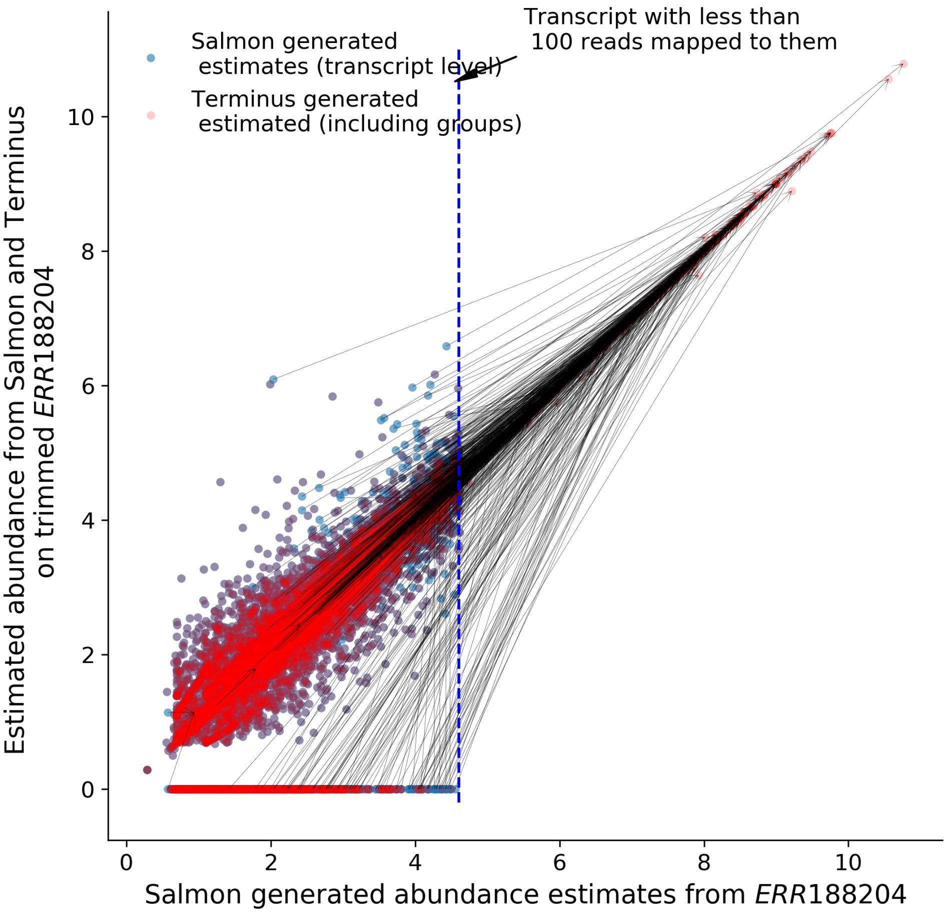


Supplementary Figure S7: The effect of groping transcript by terminus following the same convention of S6. The dashed blue vertical line signifies the transcripts which are expressed with a read count of 100.

#### 5 Biological relevance of terminus groups

The groups created from terminus are strictly data-driven, meaning the presence of uncertainty within the dataset drives the formation of specific groups. In an RNA-seq experiment it is often the case that the assigned reads are not enough to resolve the abundance values at the level of transcripts for some transcripts, while sufficient information is present to perform accurate estimation for other transcripts. In such a situation, fully relying on the higher level annotation (genes or gene families) to collapse all transcripts falling under this annotation may not be the ideal solution. Moreover, summing up transcripts at that level would eliminate transcript-level inference for the transcripts for which accurate estimation was possible, thereby defeating the purpose of transcript-level analysis. Providing an intermediate solution, terminus aims to group the transcripts for which the posterior sampling shows a high degree of uncertainty, while keeping the other transcript estimates unchanged.

To verify the biological plausibility of groups produced by terminus, we have closely-analyzed the relation between groups generated on the simulated 4 vs. 4 human dataset with the corresponding gene

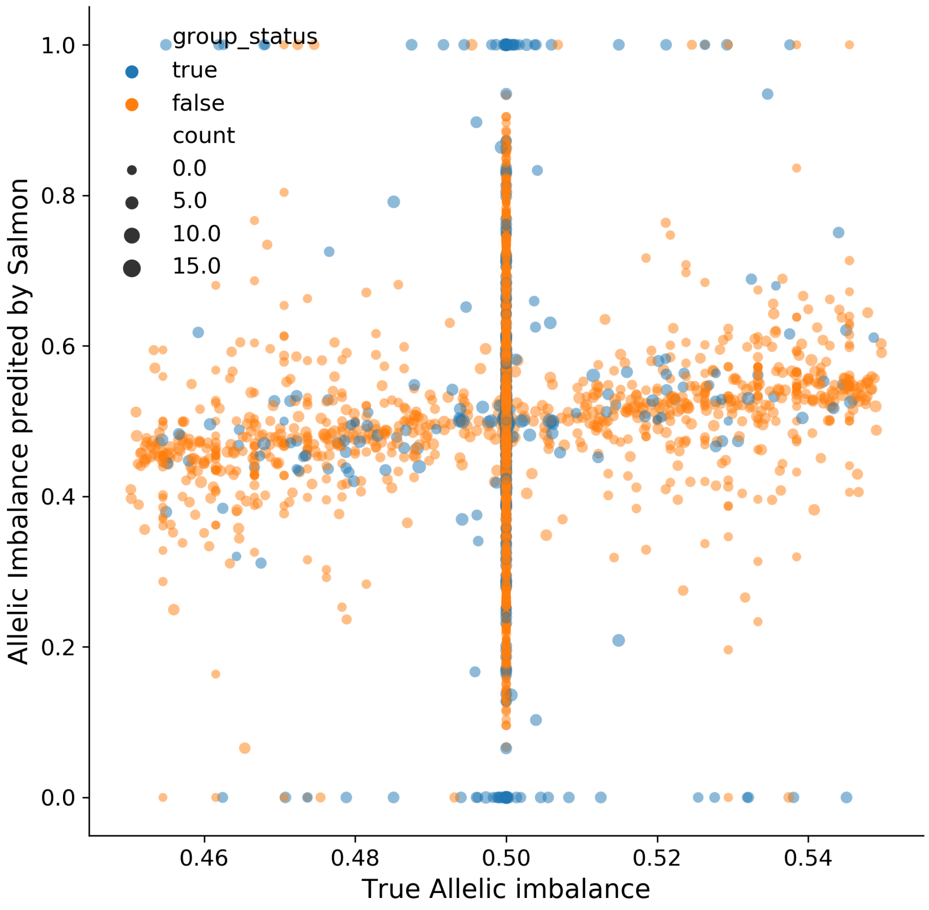


Supplementary Figure S8: Demonstration of the effect of allelic imbalance. The size of the points are determined by log of their raw counts. The horizontal line is formed by the dots that has true allelic imbalance of 0.5

families. One motivating example is the gene family *Clustered protocadherins* or clustered Pcdhs. In the present annotation[1], there are 138 transcripts that are distributed over 59 genes. Instead of grouping the entire family, terminus groups 15 genes within a group. One possible reason for such grouping by terminus is the presence of highly-ambiguous reads that are reported by *Salmon*, as shown in Supplementary Figure S9. We believe identifying such grouping within gene family may be useful for many other downstream analysis.

#### 6 Comparison with random grouping

In order to verify the efficacy of the groupings produced by terminus, we have generated a random partition within the set of transcripts following the same distribution of group sizes generated by terminus. We observe that a random grouping does not improve the accuracy from the original (ungrouped)

[1]https://biomart.genenames.org/martform/#!/default/HGNC?datasets=hgnc_family_mart

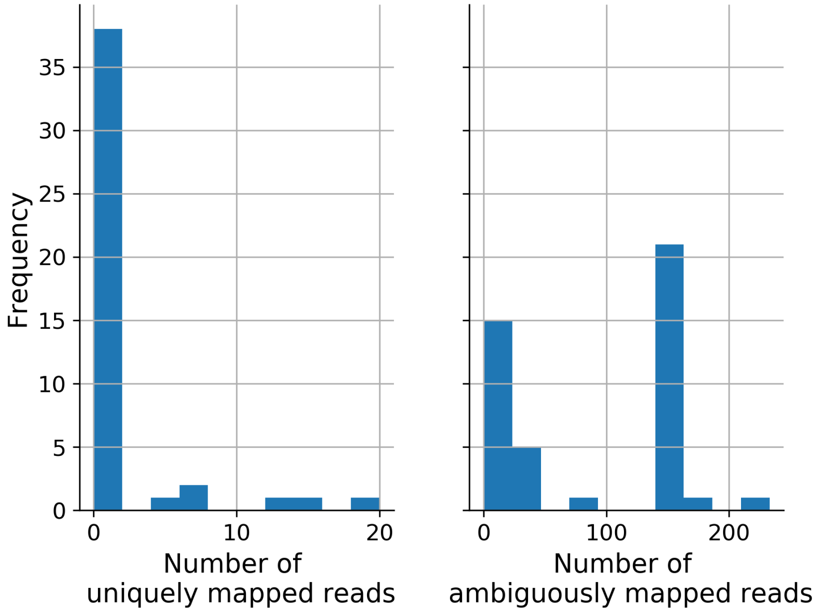


Supplementary Figure S9: Histogram of number of transcripts with respect to uniquely mapped reads and ambiguously mapped reads to from Clustered protocadherins gene family reported by *Salmon*

Supplementary Table 2: Spearman correlation and MARD for simulated 4 vs. 4 human dataset with comparing random partition vs the groups produced by respective algorithms.

| Datasets | Correlation (Spearman) | | | MARD | | |
|---|---|---|---|---|---|---|
| | *Salmon* | random | terminus | *Salmon* | random | terminus |
| simulated 4 vs. 4 human dataset | 0.94 | 0.94 | 0.96 | 0.11 | 0.12 | 0.09 |

estimates at all. Likewise, random grouping does not decrease the accuracy, as one would expect the distribution of errors over random groups to mirror the distribution of estimation errors made at the transcript-level if the grouped transcripts are not related in any meaningful way. The result on one of the simulated 4 vs. 4 human dataset samples are presented in Supplementary Table 2.

### 7 Tuning terminus to attain different number of groups

Terminus accepts several tuning parameters that can be used to control the number of groups. The most effective control on the number of groups can be achieved by using changing the consensus threshold that determines what fraction of samples should include the group, in order to count it towards the final group. For simulated 4 vs. 4 human dataset we have changed the consensus threshold parameter from 0.125 to 1.0, which dictates the number of groups when, a group has to be present in at least one sample to the condition where the group has to be present in all samples. Supplementary Table 3 shows the effect in the number of groups and corresponding correlation when we change the consensus threshold.

Supplementary Table 3: Spearman correlation and number of groups for simulated 4 vs. 4 human dataset with different values of consensus threshold

| Consensus threshold | Number of groups | Spearman Correlation |
|---|---|---|
| 0.125 | 7225 | 0.97 |
| 0.25 | 6192 | 0.96 |
| 0.50 | 4972 | 0.96 |
| 0.75 | 3861 | 0.96 |
| 1.00 | 2544 | 0.95 |

### References


Foulds, J. *et al.* (2013). Stochastic collapsed variational Bayesian inference for latent Dirichlet allocation. In *Proceedings of the 19th ACM SIGKDD international conference on Knowledge discovery and data mining*, pages 446–454.

Frazee, A. C. *et al.* (2015). Polyester: simulating RNA-seq datasets with differential transcript expression. *Bioinformatics*, **31**(17), 2778–2784.

Langmead, B. *et al.* (2009). Ultrafast and memory-efficient alignment of short DNA sequences to the human genome. *Genome biology*, **10**(3), R25.

Lappalainen, T. *et al.* (2013). Transcriptome and genome sequencing uncovers functional variation in humans. *Nature*, **501**(7468), 506–511.

Li, B. *et al.* (2010). Rna-seq gene expression estimation with read mapping uncertainty. *Bioinformatics*, **26**(4), 493–500.

Nariai, N. *et al.* (2013). TIGAR: transcript isoform abundance estimation method with gapped alignment of RNA-Seq data by variational Bayesian inference. *Bioinformatics*, **29**(18), 2292–2299.

Patro, R. *et al.* (2017). Salmon provides fast and bias-aware quantification of transcript expression. *Nature Methods*, **14**(4), 417.

Turro, E. *et al.* (2011). Haplotype and isoform specific expression estimation using multi-mapping RNA-seq reads. *Genome Biology*, **12**(2), R13.

Zakeri, M. *et al.* (2017). Improved data-driven likelihood factorizations for transcript abundance estimation. *Bioinformatics*, **33**(14), i142–i151.
